# An Efficient Algorithm for Estimating Population History from Genetic Data

**DOI:** 10.1101/2021.01.23.427922

**Authors:** Alan R. Rogers

## Abstract

The Legofit statistical package uses genetic data to estimate parameters describing population history. Previous versions used computer simulations to estimate probabilities, an approach that limited both speed and accuracy. This article describes a new deterministic algorithm, which makes Legofit faster and more accurate. The speed of this algorithm declines as model complexity increases. With very complex models, the deterministic algorithm is slower than the stochastic one. In an application to simulated data sets, the estimates produced by the deterministic and stochastic algorithms were essentially identical. Reanalysis of a human data set replicated the findings of a previous study and provided increased support for the hypotheses that (a) early modern humans contributed genes to Neanderthals, and (b) a “superarchaic” population (which separated from all other humans early in the Pleistocene) was either large or deeply subdivided.

## 1 Introduction

Legofit is a publicly-available statistical package that uses genetic data to estimate the history of size, subdivision, and gene flow within a set of populations.^1^ Because it ignores the within-population component of genetic variation, it avoids the need to estimate parameters describing recent population history and is able to focus on a deeper time scale. It operates by fitting models of history to the frequencies of “nucleotide site patterns,” which describe the sharing of derived alleles by subsets of populations. In recent publications, it has shown that Neanderthals and Denisovans separated earlier than previously thought, that their ancestors endured a bottleneck in population size, and that these ancestors interbred with a preexisting “superarchaic” population, which had inhabited Eurasia since early in the Pleistocene. It has also confirmed a variety of results first obtained by other methods [20–22].

Legofit’s estimation procedure evaluates the fit of model to data at many sets of parameter values. In previous versions of Legofit, each evaluation required a lengthy computer simulation. These calculations were feasible because they could be done in parallel. Nonetheless, Legofit was most useful on high-performance computing clusters. This stochastic algorithm also limited the accuracy with with models could be fit to data.

This article describes a new deterministic algorithm, which increases both speed and accuracy. With the simulated data discussed below, the deterministic algorithm is over 1600 times as fast as the stochastic one. Because of its greater accuracy, it also provides a better fit of model to data and improves Legofit’s ability to discriminate among models.

## 2 Methods

The new algorithm involves two novel components. The first of these involves a well-known Markov chain [8, 23, 26] that is seldom used because of the numerical difficulties. Below, section 2.3 shows a way around these difficulties. The new algorithm also relies on two results describing how descendants are partitioned among ancestors. One of these (Eqn. 7) is old and the other (Eqn. 8) new. Before discussing these, however, let us review the basics of Legofit. As in previous publications, I use capitalization to distinguish the Legofit package from the legofit program within that package.

### 2.1 Model of population history

Fig. 1 shows a gene tree embedded within a network of populations. In Legofit, the population network is modeled as a set of connected segments, each with a simple history. Each segment describes a single randomly-mating population, during an interval of constant population size. The root segment has no parent, and tip segments have no children. All other segments have at least one parent and one child. Segments that receive gene flow have two parents: one for native ancestors and the other for immigrants. Most segments have finite length, but the root segment is infinite.

**Figure 1.**
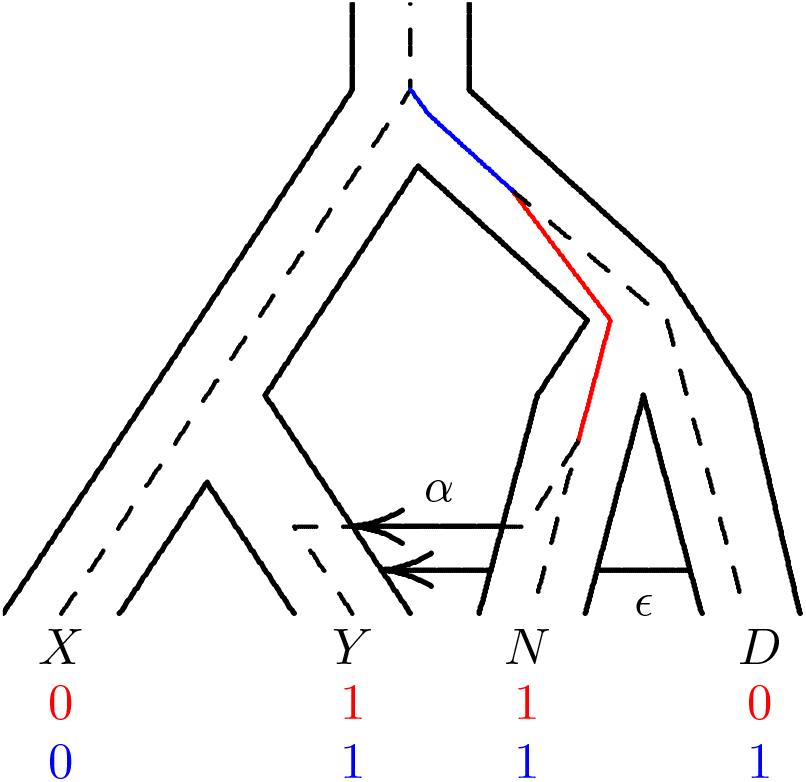
Population network with embedded gene tree. A mutation on the solid red branch would generate site pattern *yn* (shown in red at the base of the tree). One on the solid blue branch would generate *ynd*. “0” and “1” represent the ancestral and derived alleles. Key: *X*, Africa; *Y*, Eurasia; *N*, Neanderthal; *D*, Denisovan. After Rogers [20, Fig. 1].

The population history in Fig. 1 could be modeled using the network of segments in Fig. 2. Note that the branch ending at *Y* in Fig. 1 has three segments (y, y1, and y2) in Fig. 2. This is because that branch is interrupted by two episodes of gene flow, and gene flow can occur only at the ancient end of a segment. Thus, segment y extends from the present back to the first episode of gene flow, y1 extends from the first episode to the second, and y2 extends from the second episode back to the separation of populations *X* and *Y*.

**Figure 2.**
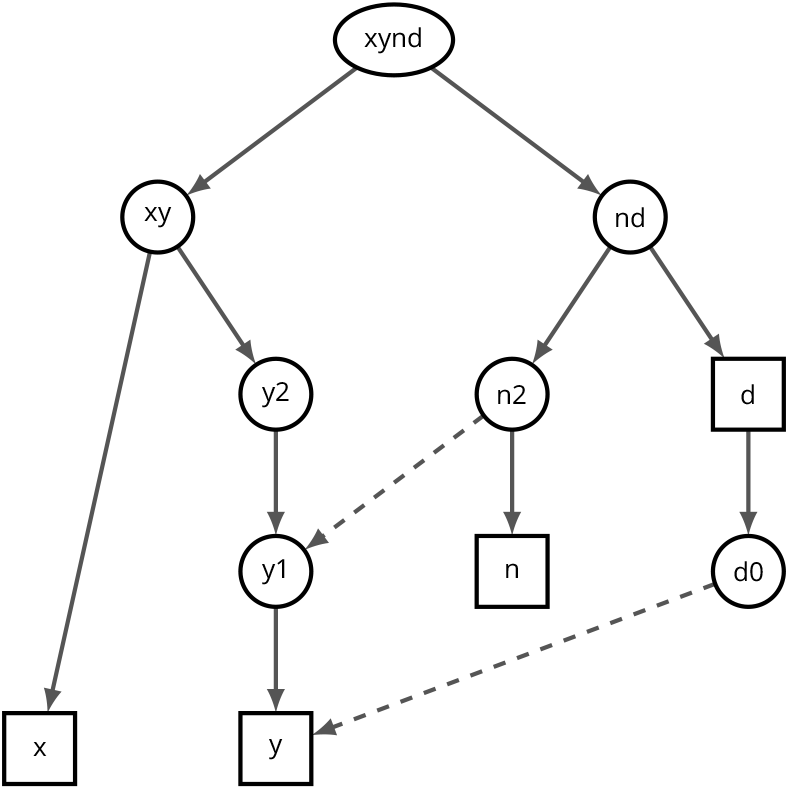
Network of segments used in legofit analysis. Squares represent segments from which we have “observed” (i.e. simulated) data. Arrows indicate ancestor-descendant relationships, and dashed lines represent gene flow. Segments in the same row need not be contemporary.

The size of population *Y* cannot be estimated, because there is never more than a single lineage within *Y*. At time zero, there is a single haploid sample, because *Y* is a population that has been sampled. This lineage may derive from segment d0, from n2, or from y2. But there is no way, under this model of history, for any of the segments that compose *Y* to contain more than one lineage. Consequently, no coalescent events are possible within *Y*, and its population size does not affect site pattern frequencies. This population size is therefore treated as a fixed constant rather than a parameter to be estimated.

On the other hand, segment n2 may contain either 1 or 2 lineages. It will always contain at least 1 lineage, which is ancestral to the lineage sampled in segment n. In addition, it may contain the lineage sampled in segment y. Consequently, population size in segment n2 is an estimable parameter.

In order to reduce the parameter count, it is possible to specify that several segments share a single population-size parameter.

### 2.2 Nucleotide site patterns

Legofit works with the frequencies of *nucleotide site patterns*, which are illustrated in Fig. 1. A nucleotide site exhibits the *yn* site pattern if random nucleotides drawn from populations *Y* and *N* carry the derived allele, but those drawn from other populations carry the ancestral allele. Fig. 1 shows the gene genealogy of a particular nucleotide site, embedded within the network of populations. A mutation on the red branch would generate site pattern *yn*, whereas one on the blue branch would generate *ynd*. Mutations elsewhere would generate other site patterns. The gene genealogy will vary from locus to locus, so averaging across the genome involves averaging across gene genealogies. We are interested in the properties of such averages.

Let *B*_*i*_ represent the length in generations of the branch generating site pattern *i*. I employ the “infinite sites” model of mutation [10], which assumes that the mutation rate is small enough that we can ignore the possibility of multiple mutations on any given branch. Under this assumption, a polymorphic site exhibits pattern *i* with probability

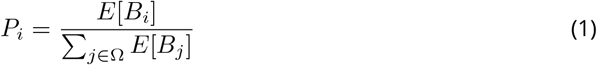

where *E*[*B*_*i*_] is the expected length of the branch generating site pattern *i*, and Ω is the set of site patterns under study [20, Eqn. 1]. Previous versions of Legofit used coalescent simulations to estimate these expectations. The sections that follow describe a deterministic algorithm.

### 2.3 The matrix coalescent

The new algorithm is based on a model that calculates the probability that there are *k* ancestral lineages at the ancient end of a segment, given that there are *n* descendant lineages at the recent end. This model also calculates the expected length of the interval within the segment during which there are *k* lineages, where 1 ≤ *k* ≤ *n*. The model employs a continuous-time Markov chain, which begins with *n* haploid lineages at the recent end of the segment. As we trace the ancestry of this sample into the past, the original sample of *n* lineages falls to *n* − 1, then *n* − 2, and so on until only a single lineage is left, or we reach the end of the segment.

The number, *n*, of descendants equals 1 for tip segments. For ancestral segments, *n* may take several values with different probabilities. The legofit program sums across these possibilities, weighting by probability.

This Markov chain is well known [23, appendix I; 8; 26] but seldom used, because accurate calculations are difficult with samples of even modest size. Legofit, however, is designed for use with small samples. Furthermore, it is possible (as shown below) to factor the calculations into two steps, one of which can be done in exact arithmetic and only needs to be done once at the beginning of the computer program. Numerical error arises only in the second step, and as we shall see, that error is small.

Within a segment, the population has constant haploid size 2*N*, although 2*N* can vary among segments. (“Haploid” population size is twice the number of diploid individuals.) It will be convenient to measure time backwards from the recent end of each segment in units of 2*N* generations. On this scale, time is *ν* = *t/*2*N*, where *t* is time in generations. Let **x**(*ν*) denote the column vector whose *i*th entry, *x*_*i*_(*ν*), is the probability of observing *i* lineages at time *ν*, where 1 ≤ *i* ≤ *n*. I ignore the absorbing state *x*_1_, so that indices of arrays and matrices range from 2 to *n*. Because there are *n* lineages at time zero (the recent end of the segment), the initial vector equals **x**(0) = [0, …, 0, 1]^*T*^. At time *ν* [26, Eqn. 8],

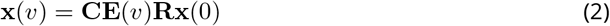

Here, **E**(*ν*) is a diagonal matrix of eigenvalues whose *i*th diagonal entry is 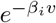, where *β*_*i*_ = *i*(*i* − 1)*/*2. **C** = [*c*_*ij*_] and **R** = [*r*_*ij*_] are matrices of column eigenvectors and row eigenvectors, both of which are upper triangular. They are calculated by setting diagonal entries equal to unity, and then applying [26, p. 1642],

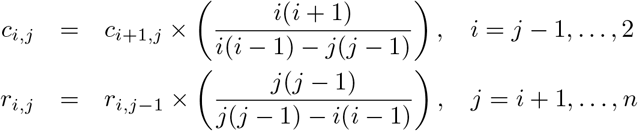

Let **m**(*ν*) denote the vector whose *k*th entry, *m*_*k*_(*ν*), is the expected duration (in units of 2*N* generations) of the interval during which the segment contains *k* lineages, within a segment of length *ν*. This vector equals

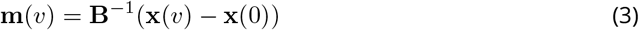

where

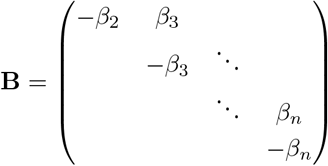

Eqn. 3 holds not only for finite segments, but also when *ν* → ∞. In the infinite case, **x**(∞) = 0, because we are considering only the transient states (*x*_2_, …, *x*_*n*_), which disappear in the long run. Eqn. 3 is easy to calculate, because **B**^−1^ has a simple form. For the case of *n* = 4,

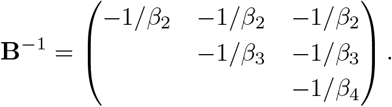

This model presents challenging numerical issues. To deal with these, let us re-organize the calculations to do as much as possible in exact arithmetic. I illustrate this re-organization using the case of *n* = 3, for which Eqn. 2 becomes

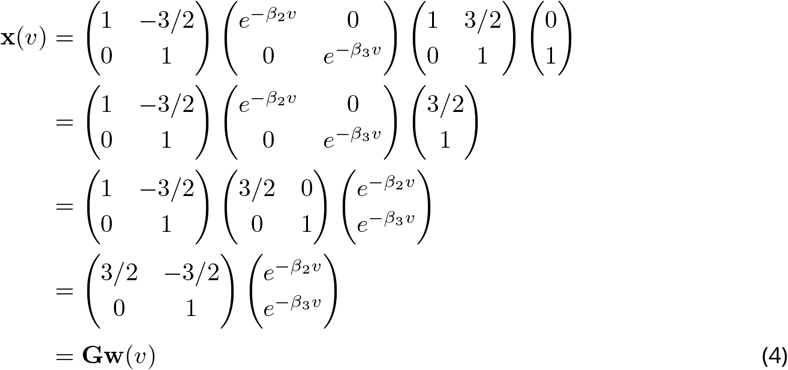

where 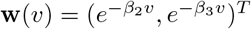 is a vector of eigenvalues, **G** = **C** diag(**Rx**) is a matrix of column eigenvectors with columns scaled by the entries of vector **Rx**(0), and diag(**Rx**(0)) is a diagonal matrix whose main diagonal equals the vector **Rx**(0). The matrix **G** can be calculated in exact rational arithmetic. This is done at the beginning of the computer program for each possible value of *n*, and the resulting values are stored for later use.

Next, substitute (4) into (3) to obtain

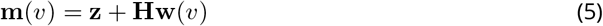

where **z** = −**B**^−1^**x**(0) = (1*/β*_2_, …, 1*/β*_*n*_)^*T*^, and **H** = **B**^−1^**G**, both of which can be calculated in advance for each possible value of *n*, using exact arithmetic. For example, if *n* = 3,

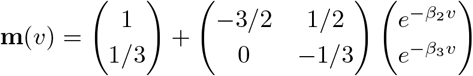

In an infinite segment, Eqn. 5 is simply **m**(∞) = **z**.

This algorithm calculates *x*_*k*_(*ν*) and *m*_*k*_(*ν*) only for *k* = 2, 3, …, *n*. Values for *k* = 1 are obtained by subtraction: 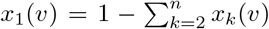, and 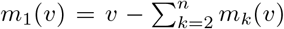. Finally, to re-express *m*_*k*_(*ν*) in units of generations, define

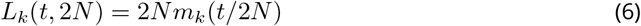

where *t* is the length of the current segment in generations, and 2*N* is its haploid population size. *L*_*k*_(*t*, 2*N*) is the expected duration in generations of the interval during which the current segment contains *k* lineages.

Several of the quantities in this algorithm—**G, H**, and **z**—are calculated in exact rational arithmetic. Although there is no roundoff error, these calculations will overflow if *n* is too large. With 32-bit signed integers, there is no overflow until *n >* 35. This is more than enough for Legofit, which requires that *n* ≤ 32, so that site patterns can be represented by the bits of a 32-bit integer.

Roundoff error does occur in this algorithm, because all quantities are eventually converted to double-precision floating point during the calculation of Eqns. 4 and 5. To assess the magnitude of this error, I compared results to calculations done in 256-bit floating-point arithmetic, using the Gnu MPFR library [7]. I considered values of *ν* ranging from 0 to 9.5 in steps of 0.5, and also *ν* → ∞. The maximum absolute error is 3.553 *×* 10^−15^ when *n* = 8; 2.700 *×* 10^−13^ when *n* = 16; and 1.543 *×* 10^−8^ when *n* = 32. These errors are all much smaller than those of Legofit’s stochastic algorithm.

The theory just described allows us to calculate the probability that *n* descendants have *k* ≤ *n* ancestors in some previous generation. To relate this theory to the frequencies of site patterns, we must discuss how the coalescent process partitions descendants among ancestors.

### 2.4 Partitioning descendants among ancestors

A “segment” is an interval within the history of one subpopulation. Let *n* represent the number of descendant lineages at the recent end of the segment, and let *k* ≤ *n* represent the number of ancestral lineages at some earlier point within the segment. The theory in section 2.3 calculates the probability of *k* at any time within the segment and also provides the expected length of the subinterval containing *k* lines of descent.

For all segments except the root, we need both of these quantities. We need the expected lengths of subintervals, because these lengths measure the opportunity for mutation. In addition, we need to assign a probability to each of the ways in which the set of descendants can be partitioned among ancestors at the ancient end of the segment. These partitions and probabilities are used in calculations on earlier segments within the network.

For the root segment, we still need the expected lengths of subintervals. But because there are no earlier segments to worry about, we don’t need to assign probabilities to partitions. This is fortunate, because the number of set partitions increases rapidly with the size of the set [11, p. 418], and the set of descendants is largest in the root segment.

To address these needs, I present two algorithms. One sums across partitions of the set of descendants and is used in all segments except the root. The other avoids this sum and is used only at the root.

#### 2.4.1 Summing across set partitions

Section 2.3 calculated the expected length of the interval during which there are *k* ancestors, given that there are *n* descendants at the recent end of the segment. If a mutation strikes one ancestor, it will be shared by all descendants of that ancestor. The subset comprising these descendants corresponds to a nucleotide site pattern.

Suppose that at some time in the past there were *k* ancestors. These ancestors partition the set of descendants into *k* subsets. Let *x*_1_, *x*_2_, …, *x*_*k*_ denote the sizes of the *k* subsets, i.e., the numbers of descendants of the *k* ancestors. The conditional probability, given *k*, of such a partition is [3, theorem 1.5, p. 11]

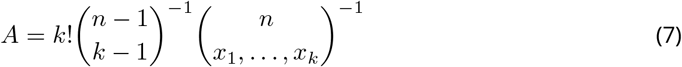

The left side of table 1 shows all ways of partitioning a set of 4 descendants among 2 ancestors along with the probability of each partition. The descendants of each ancestor define a nucleotide site pattern. For example, the first partition is “1112,” which says that the first three descendants share a single ancestor. A mutation in this ancestor would be shared by these descendants, and so the descendants correspond to a site pattern.

**Table 1.**
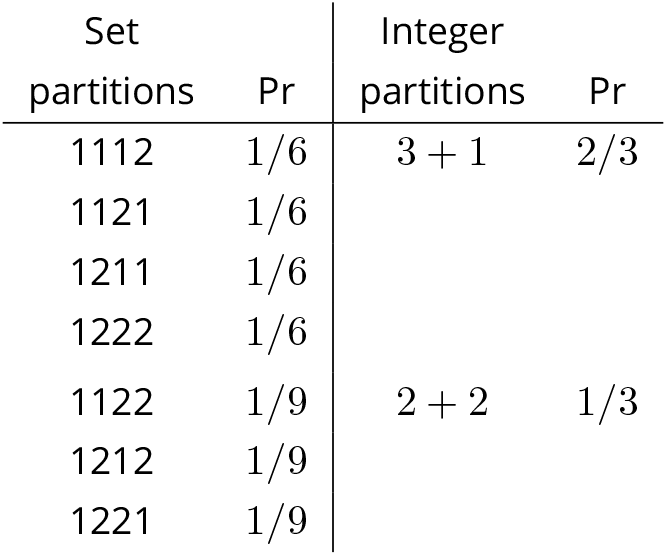
Set partitions, integer partitions, and their probabilities, for the case in which *n* = 4 and *k* = 2. Under “set partitions,” the value in position *j* of each string is the index of the ancestor of descendant *j*. Thus, “1122” means that descendants 1 and 2 descend from one ancestor, whereas 3 and 4 descend from another. Ancestors are numbered in order of their appearance in the list of descendants. Integer partitions are discussed in section A.2 of the appendix.

| Set partitions | Pr | Integer partitions | Pr |
| --- | --- | --- | --- |
| 1112 | 1/6 | 3 + 1 | 2/3 |
| 1121 | 1/6 |  |  |
| 1211 | 1/6 |  |  |
| 1222 | 1/6 |  |  |
| 1122 | 1/9 | 2 + 2 | 1/3 |
| 1212 | 1/9 |  |  |
| 1221 | 1/9 |  |  |

This result is used in an algorithm that calculates (a) all possible partitions of descendants at the ancient end of the segment along with their probabilities, and (b) the contribution of the current segment to the expected branch length of each site pattern. The algorithm loops first across values of *k*, where 1 ≤ *k* ≤ *n*. For each *k*, it loops across set partititions using Ruskey’s algorithm [11, pp. 764–765]. The probability that a given partition occurs at the ancient end of a segment, given the set of descendants at its recent end, is the product of *x*_*k*_(*t/*2*N*) (Eqn. 2) and *A* (Eqn. 7). Each partition also makes a contribution to the expected branch length associated with *k* site patterns—one for each ancestor. That contribution is the product of *L*_*k*_(*t*, 2*N*) (Eqn. 6) and *A* (Eqn. 7). These contributions are summed across partitions and segments to obtain the expected branch length of each site pattern.

#### 2.4.2 A faster algorithm for the root segment

Consider the event that a particular set of *d* descendants (and no others) descend from a single ancestor in some previous generation, given that there were *k* ancestors in that generation. This event is of interest, because a mutation in this ancestor would be shared uniquely by the *d* descendants. The probability of this event is

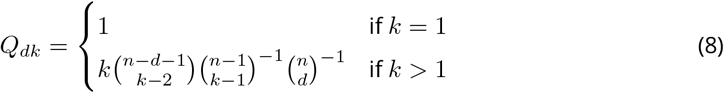

To justify this result, consider first the case in which *k* = 1. This requires that all *n* descendants descend from a single ancestor, so *d* must equal *n*. There is only one way this can happen, and because the probability distribution must sum to 1, it follows that *Q*_*dk*_ = 1. The result for *k >* 1 is derived in appendix A.

##### Example 1

Suppose *k* = *n*. In this case, each ancestor has 1 descendant, so *d* = 1, and *Q*_1,*n*_ must equal 1. Equation 8 agrees:

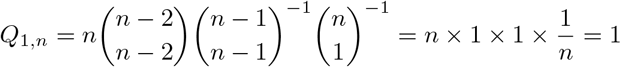

##### Example 2

Suppose that *k* = *n* − 1. In this case, we are reckoning descent from the previous coalescent interval, in which there were *n* − 1 ancestors. Consider first the case in which *d* = 1. Among the *n* descendants, 2 derive from an ancestor that split, and *n* − 2 derive from one that did not split. This implies that *Q*_1,*n*−1_ equals (*n* − 2)*/n*, the probability a random descendant derives from an ancestor that did not split.

The case of *d* = 2 is also easy. There are 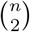 ways to choose 2 descendants from *n*, and only one of these pairs derives from a single ancestor in the previous coalescent interval. Thus, 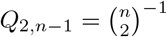. Equation 8 confirms both of these results:

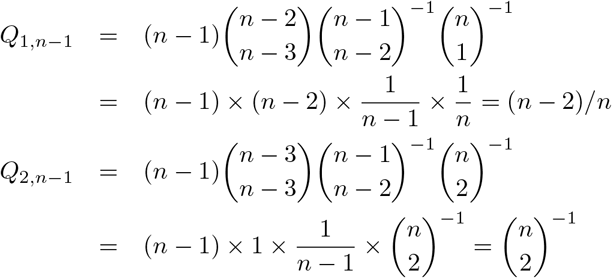

##### Example 3

We can also evaluate Eqn. 8 by comparing its results to Eqn. 7. Table 1 shows all partitions and their probabilities for the case in which *k* = 2 and *n* = 4. Notice that subsets of sizes 1, 2, and 3 have probabilities 1/6, 1/9, and 1/6. Eqn. 8 yields identical values:

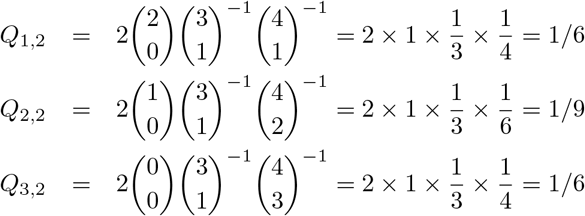

In the root segment, the program uses the following algorithm: Loop first across values of *k*, where 1 ≤ *k* ≤ *n*. For each *k*, loop across values of *d*. If *k* = 1, then *d* = *n*. Otherwise, *d* can take any integer value such that 1 ≤ *d* ≤ *n* − *k* + 1. For each *d*, calculate *Q*_*dk*_ using Eqn. 8, and loop across ways of choosing *d* of *n* descendants, using algorithm T of Knuth [11, p. 359]. Each such choice corresponds to a nucleotide site pattern. Add *Q*_*dk*_*L*_*k*_(*t*, 2*N*) to the expected branch length associated with this site pattern.

### 2.5 Simulated data sets

To evaluate the new algorithm, I used 50 data sets simulated with msprime [9], using the model in Fig. 1, which is identical to that used in a previous publication [20]. Each simulated genome consisted of 1000 chromosomes, each with 2 *×* 10^6^ nucleotide sites. Each simulated data set consisted of 4 genomes, one each from populations *X, Y, N*, and *D*, which represent the African, European, Neanderthal, and Denisovan populations. *XY* is the population ancestral to *X* and *Y, ND* is that ancestral to *N* and *D*, and *XY ND* is that ancestral to *X, Y, N*, and *D*. The mutation rate was 1.4 *×* 10^−8^ per base pair per generation, and the recombination rate was 10^−8^ per base pair per generation.

The time parameters in the simulation model, expressed in generations, are as follows:

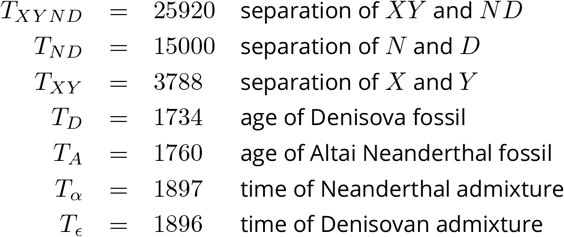

Admixture proportions are:

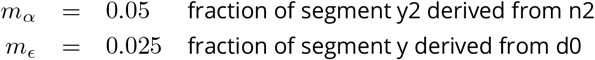

Population sizes are expressed as “haploid” counts, which represent twice the number of diploid individuals. These parameters are:

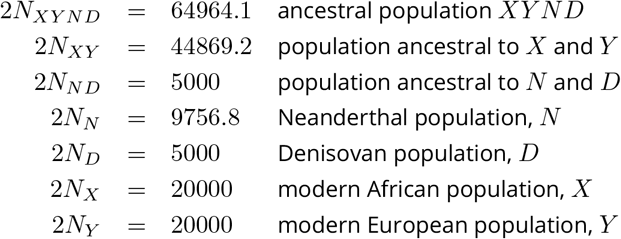

Simulation code is in section S1 of Supplementary Materials. Simulation results are in the archive (doi:10.17605/OSF.IO/74BJF).

### 2.6 Analysis of simulated data

The data analysis pipelines for the deterministic and stochastic algorithms are detailed in supplementary section S2. In both cases, the analysis was based on a model of history specified by the input file a.lgo (supplementary section SB.1). This file defines the network of segments shown in Fig. 2.

Several of the parameters of the simulation model were treated as fixed constants, because their values have no effect on expected site pattern frequencies: 2*N*_*X*_, 2*N*_*Y*_, *T*_*α*_, and *T*_ϵ_. Another parameter, *T*_*XY ND*_, was fixed at its true value to calibrate the molecular clock. The remaining 11 parameters were estimated.

For both algorithms, data analysis involved 5 stages. In stage 1, legofit was run on each of 50 simulated data sets. Each run produced two output files: a .legofit file, which contains parameter estimates, and a .state file, which records the state of the optimizer at the end of the run. The optimizer uses the *differential evolution* algorithm [17]. This algorithm maintains a swarm of points, each of which represents a guess about the values of the free parameters. There are ten times as many points as free parameters, as recommended by Price et al. [17].

Although differential evolution is good at finding global optima, it is possible that some of the stage 1 runs will get stuck on different local optima. Stage 2 is designed to avoid this problem. Each job in stage 2 begins by reading all 50 of the .state files produced in stage 1, and sampling among these to construct a swarm of points. This allows legofit to choose among local optima.

Figure 3 plots pairs of free parameters after stage 2 of the analysis. Each sub-plot has 50 points, one for each simulated data set. Several pairs of parameters are tightly correlated, and these correlations reflect “identifiability” problems: different sets of parameter values imply almost identical site pattern frequencies. To ameliorate this problem, stage 3 of the analysis uses the *pclgo* program to perform a principal components analysis, which re-expresses the free variables in terms of uncorrelated principal components (PCs). In previous publications [20–22], we used this step to reduce the dimension of the analysis, by excluding components that explain little of the variance. However, excluding dimensions can introduce bias, especially in the presence of identifiability problems, so I chose here to retain the full dimension. Even without any reduction in dimension, re-expression in terms of PCs improves the fit of model to data, because it allows legofit to operate on uncorrelated dimensions.

**Figure 3.**
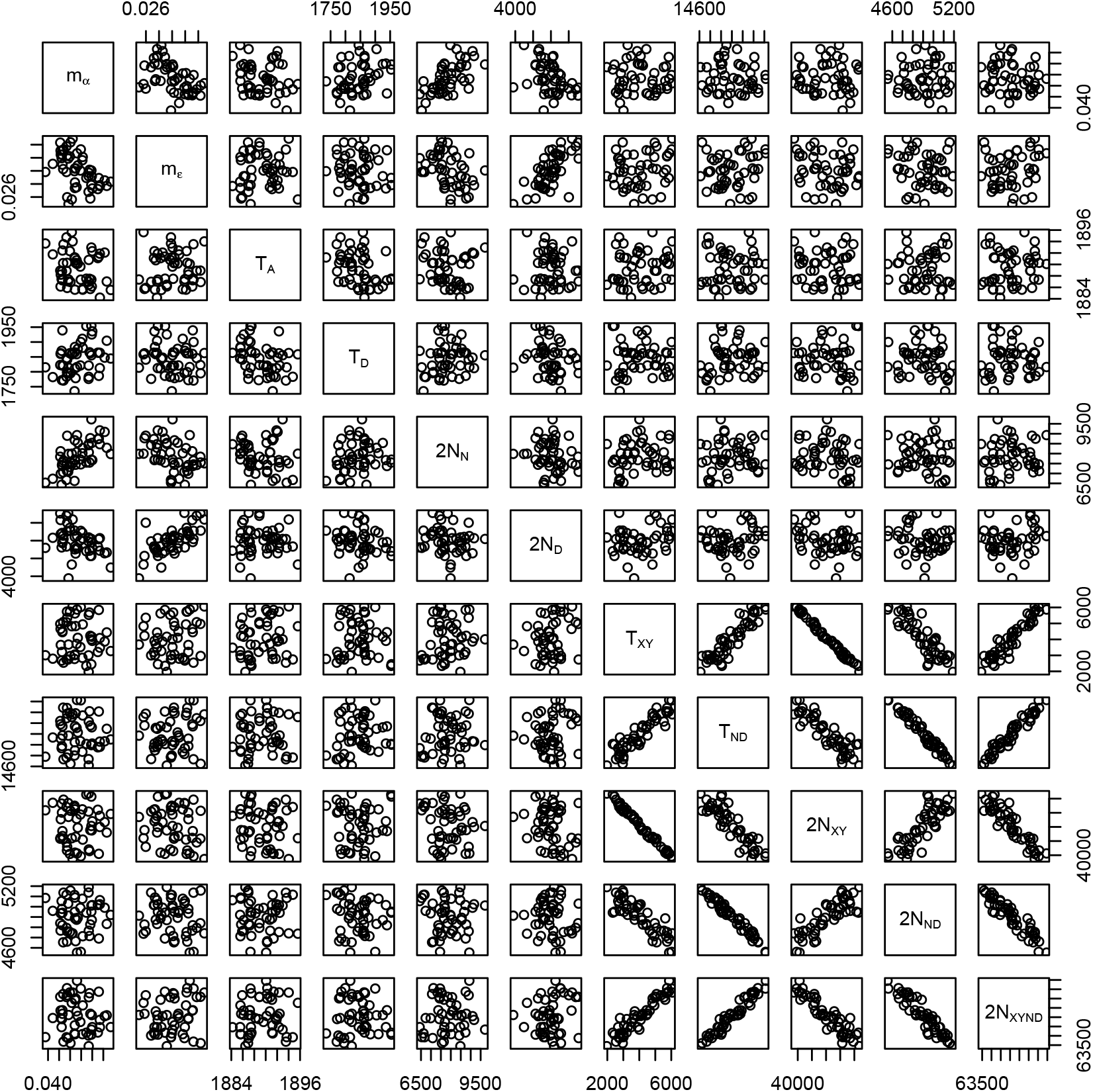
Scatter plot of each parameter against each other, based on 50 simulated data sets.

Stages 4 and 5 are like stages 1 and 2, except that the free variables are re-expressed in terms of PCs.

The program uses KL divergence [13] to measure the discrepancy between observed and predicted site pattern frequencies. Minimizing KL divergence is equivalent to maximizing multinomial composite likelihood. The optimizer stops after a fixed number of iterations or when the difference between the best and worst KL divergences falls to a pre-determined threshold. This threshold was 3 *×* 10^−6^ for the deterministic algorithm and 2 *×* 10^−5^ for the stochastic algorithm. This difference reflects the fact that the deterministic algorithm is capable of much greater precision.

### 2.7 Analysis of speed as a function of model complexity

As model complexity increases, the number of states increases. This reduces the speed of the deterministic algorithm and increases memory usage. To study this effect, I used the *legosim* program, which calculates the site pattern frequencies implied by a given model. I studied a series of models without migration or changes in population size. The models differed in the number of populations, which ranged from four to nine. Timings were done on a 2018 MacBook Air.

### 2.8 Analysis of real data

I used the deterministic algorithm to replicate the analysis of Rogers et al. [22]. (Data sets and analysis files are in directory xyvad of the archive (doi:10.17605/OSF.IO/74BJF).) That paper studied modern human sequence data from Europe and Africa [15], along with three high-coverage archaic genomes: two Neanderthals (Altai [19] and Vindija [18]), and one Denisovan [16]. It analyzed these data under eight different models, all of which are based on the history in Fig. 4.

**Figure 4.**
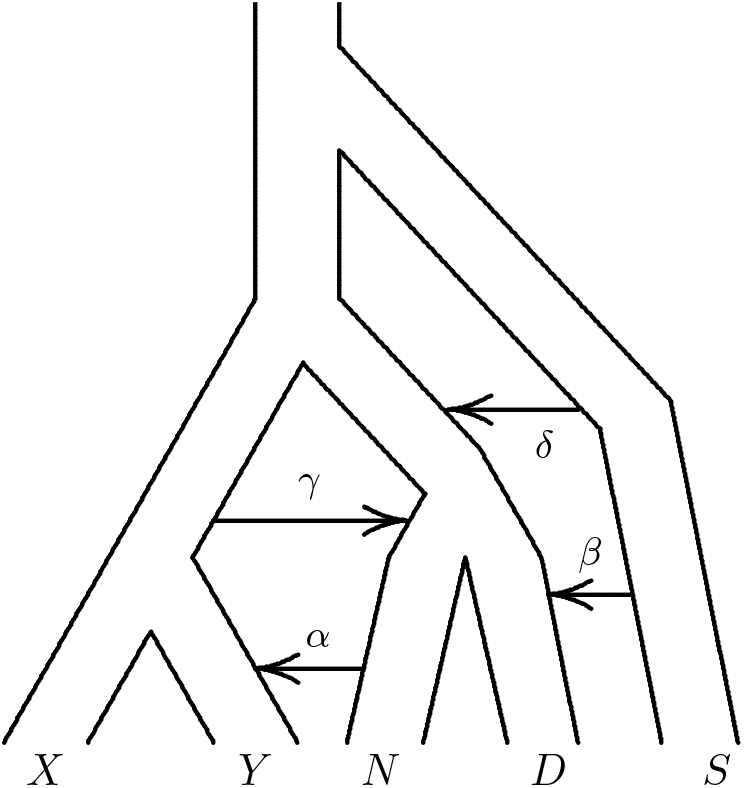
A population network including four episodes of gene flow. Upper case letters (*X, Y, N, D*, and *S*) represent populations (Africa, Europe, Neanderthal, Denisovan, and superarchaic). Greek letters label episodes of admixture.

In that figure, capital roman letters refer to populations: *X* is Africa, *Y* is Europe, *N* is Neanderthal, *D* is Denisovan, and *S* (for “superarchaic” [19]) is a population that separated from all other humans early in the Pleistocene. Greek letters label episodes of admixture. Episode *α* refers to admixture from Neanderthals into Europeans, *β* to admixture from superarchaics into Denisovans [12, 18, 19, 24, 25], *γ* to admixture from early moderns into Neanderthals [12], and *δ* to admixture from superarchaics into the “neandersovan” ancestors of Neanderthals and Denisovans [22].

Following Rogers et al. [22], I considered eight models, all of which include *α*, and including all combinations of *β, γ*, and/or *δ*. I label models by concatenating Greek letters. For example, *αβ* is the model that includes *α* and *β* but not *γ* and *δ*. This analysis is described in section S3 of the supplement.

## 3 Results and Discussion

I used both algorithms—one deterministic and the other stochastic—to fit 50 simulated data sets. In each case, this involved 200 runs of the legofit program—4 for each of 50 data sets—and 1 run of pclgo. Altogether, the deterministic version of this analysis took 18.7 CPU minutes. Because these calculations were parallelized, the elapsed time was only 1.7 minutes. Using the stochastic algorithm, the same analysis took 514.8 CPU hours, or 11.4 hours of elapsed time. For this model, the deterministic algorithm is 1654 times as fast as the stochastic one.

These timings were done on a node at the Center for High Performance Computing (CHPC) at the University of Utah, using 96 parallel threads of execution. To get a sense of how long these calculations would take on a less powerful computer, I did one run of legofit on a 2018 MacBook Air, using the deterministic algorithm with 2 threads. That took 26.2 seconds of CPU time or 13.7 seconds of elapsed time. By comparison, the CHPC node did this job in 12.4 seconds of CPU time, or 1 second of elapsed time. The high-performance node is nearly 14 times as fast as the MacBook Air, implying that the full analysis would take 24 minutes on the MacBook Air. Thus, the deterministic algorithm makes Legofit feasible on small computers.

Figure 5 shows the residual error in site pattern frequencies under the two algorithms. Residuals are substantially smaller under the deterministic algorithm because of its greater accuracy. When parameters are estimated by computer simulation, each additional decimal digit of precision requires a 100-fold increase in the number of iterations. This imposes a limit on the accuracy of the stochastic algorithm, even with the fastest computers.

**Figure 5.**
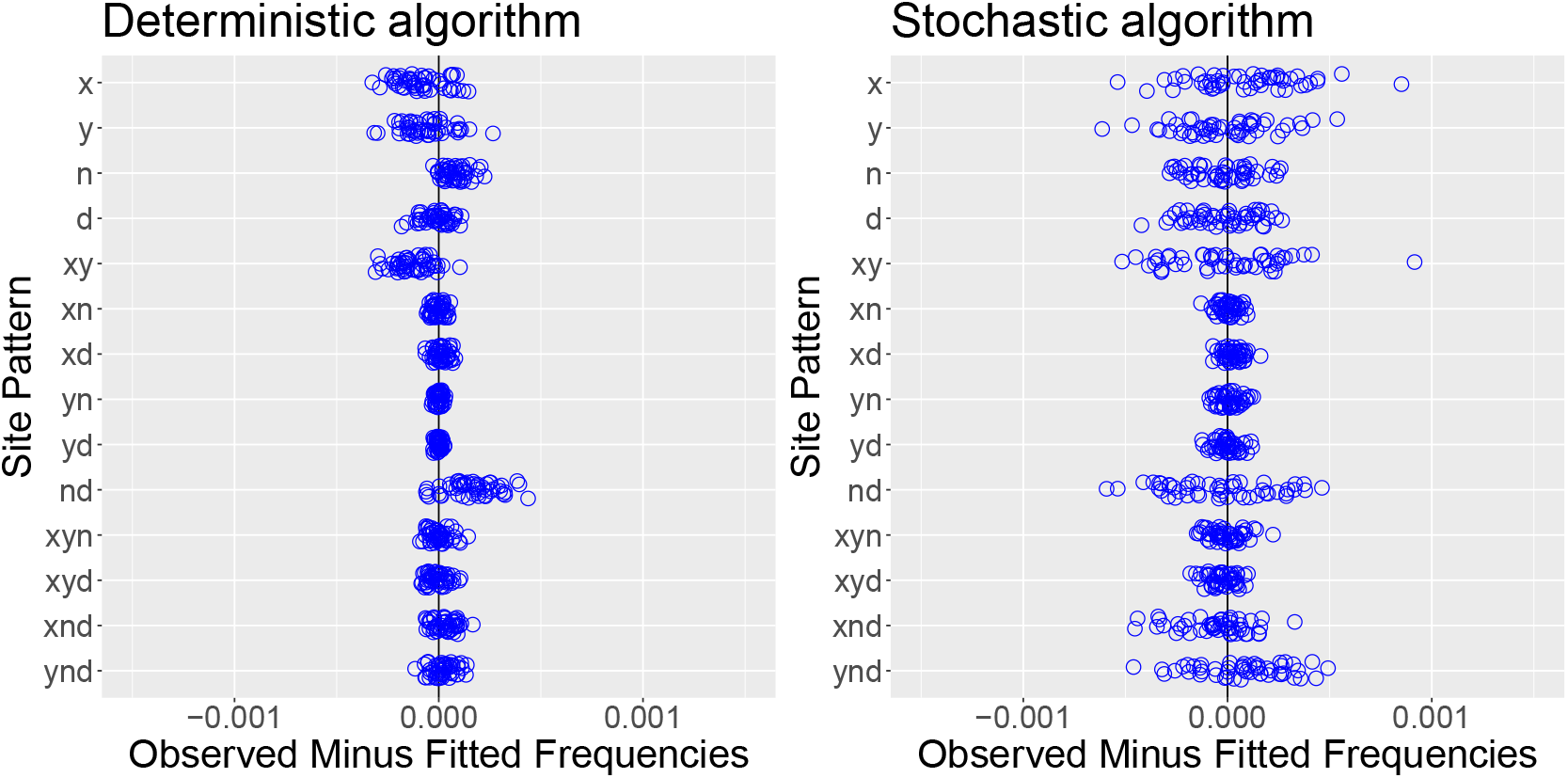
Residual error of deterministic and stochastic algorithms, based on 50 simulated data sets. Each circle refers to a different simulated data set.

To estimate site pattern frequencies, both algorithms integrate over the states of the stochastic process. The number of states increases with model complexity, so both algorithms are slower when the model is complex. Figure 6 illustrates the effect on speed. In complex models, the stochastic algorithm is faster than the deterministic one.

**Figure 6.**
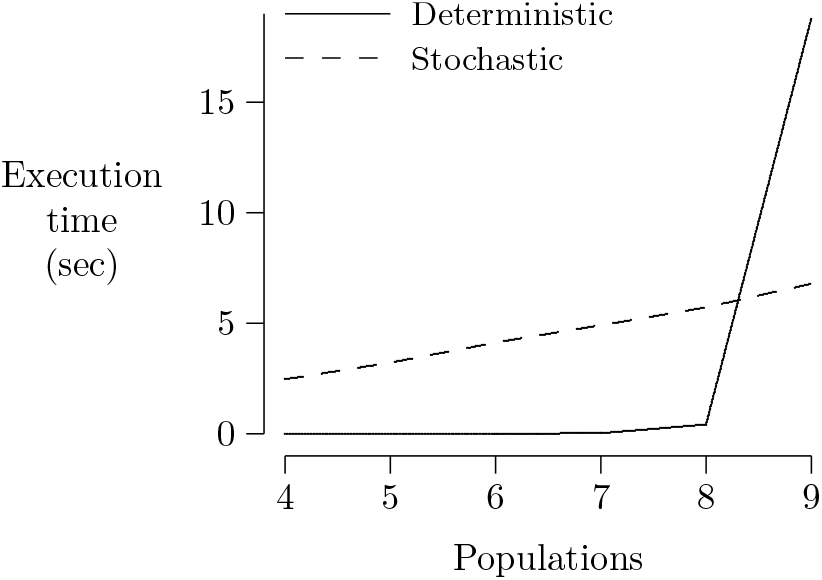
Execution time of legosim, excluding system calls, in models without migration. For the stochastic algorithm, each run used two million iterations.

Figure 7 shows the parameter estimates from the 50 data sets (blue dots) along with the true parameter values (red crosses). The two algorithms behave similarly. It does not appear that the smaller residual error of the deterministic algorithm (Fig. 5) translates into more accurate parameter estimates. This is probably because most of the spread in the parameter estimates reflects the identifiability problems seen in Fig. 3. To understand this effect, note the tight correlation between *T*_*XY*_ and 2*N*_*XY*_ in Fig. 3. This correlation exists because it is hard to distinguish the case in which 2*N*_*XY*_ is large and *T*_*XY*_ small from that in which the opposite is true. Because of this ambiguity, both parameters exhibit large uncertainties in Fig. 7.

**Figure 7.**
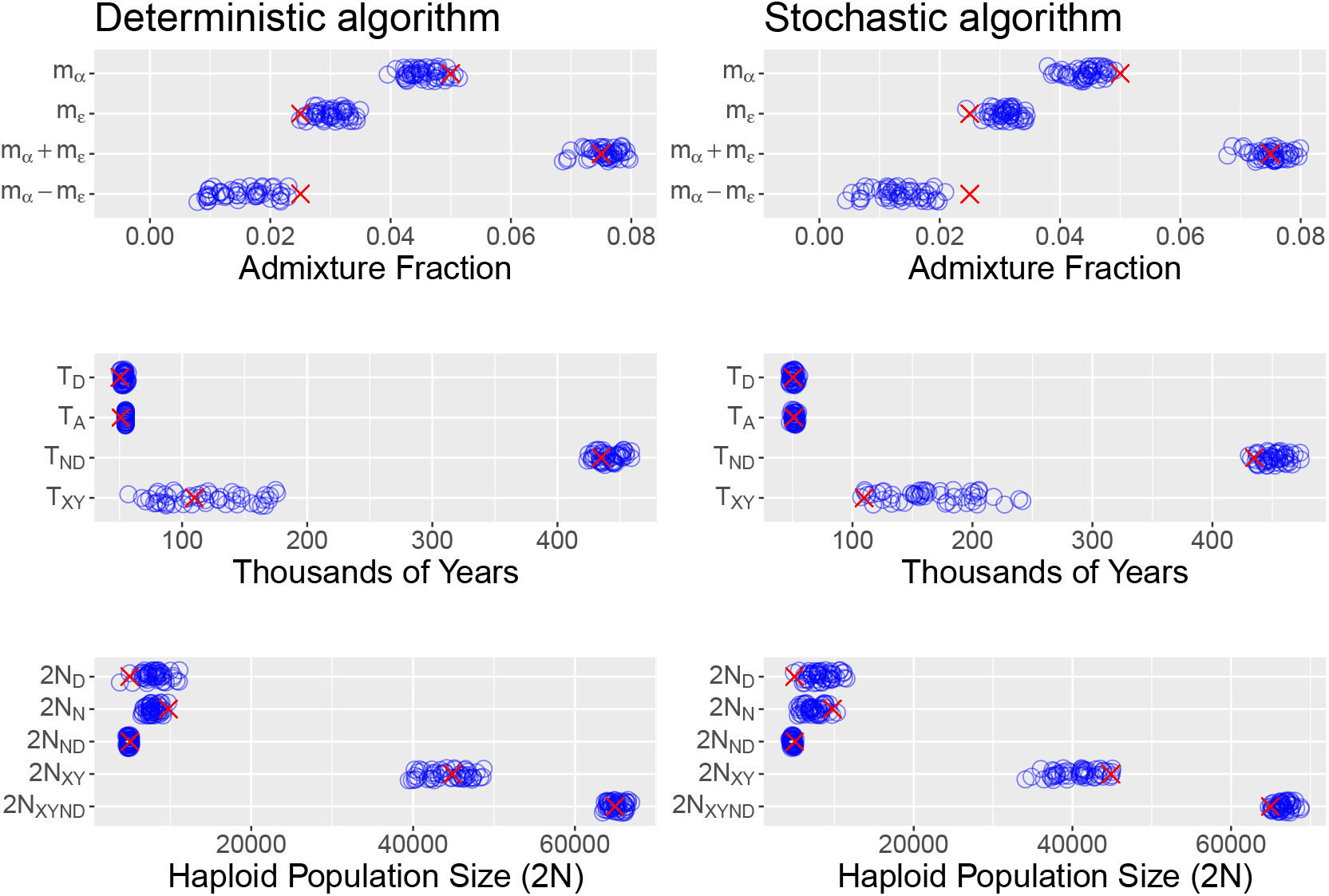
Parameter estimates from 50 simulated data sets, using the deterministic and stochastic algorithms. Blue circles are estimates and red crosses are the true parameter values.

Some bias is evident in these estimates. For example, the estimates of *m*_*α*_ tend to be a little low and those of *m*_*ϵ*_ a little high [20]. This reflects the negative correlation between these parameters that can be seen in Fig. 3. Because the two source populations (*N* and *D*) are so similar, they are hard to distinguish. We get a better estimate of the sum (*m*_*α*_ + *m* _*ϵ*_) than of the difference (*m*_*α*_ − *m*_*ϵ*_). There is also some bias in 2*N*_*D*_ and 2*N*_*N*_. In spite of these biases, the swarms of estimates tend to enclose the true parameter values, so the biases in these estimates are modest compared with their uncertainties. It should not, however, be assumed that this will always be the case.

To illustrate the new algorithm in a full-scale analsis of real data, I replicated the analysis of Rogers et al. [22]. Table 2 shows the CPU time used by each algorithm in analysis of the eight models in that publication. For this set of models, the deterministic algorithm is always faster, but its execution time ranges across several orders of magnitude. These execution times are not strictly comparable, because they involve several compute clusters, which vary in processor speed. These differences are minor, however, compared with the enormous differences in run time seen in table 2.

**Table 2.** CPU time expended in analysis of each model from Rogers et al. [22]. Each analysis involves 204 runs of legofit and 1 run of pclgo. Elapsed times were much shorter, because calculations were done in parallel. “Acceleration” is the ratio of execution speed in the deterministic model to that in the stochastic model. Models are arranged in order of increasing execution time with the deterministic algorithm.

| Model | $\log_{10}$ seconds | | Acceleration |
| --- | --- | --- | --- |
|  | Deterministic | Stochastic |  |
| $\alpha$ | 1.60246 | 5.94980 | 22250.5 |
| $\alpha\beta$ | 2.60384 | 5.81015 | 1608.1 |
| $\alpha\gamma$ | 2.83806 | 5.94417 | 1276.8 |
| $\alpha\beta\gamma$ | 3.67736 | 6.03901 | 230.0 |
| $\alpha\delta$ | 4.42860 | 6.16730 | 54.8 |
| $\alpha\beta\delta$ | 4.86285 | 6.04239 | 15.1 |
| $\alpha\gamma\delta$ | 5.47544 | 6.14022 | 4.6 |
| $\alpha\beta\gamma\delta$ | 6.04505 | 6.20171 | 1.4 |

To choose among models, I used the bootstrap estimate of predictive error, “bepe” [4, 5, 20]. This method uses variation among data sets (the real data plus 50 replicates generated by a moving-blocks bootstrap [14]) to approximate variation in repeated sampling. It fits the model to one data set and then tests this fit against all the others. Table 3 uses all models to compare the bepe values calculated by the deterministic and stochastic algorithm. In all cases, the deter-ministic algorithm yields a smaller bepe value than the stochastic algorithm, indicating a better fit of model to data. The order of the eight models, however, is the same. Because the deterministic algorithm yields smaller bepe values, one should use the same algorithm (stochastic or deterministic) for all models in any analysis. Otherwise, model selection will be biased in favor of deterministic results because of their smaller bepe values.

**Table 3.** Bootstrap estimate of predictive error (bepe) values and bootstrap model average (booma) weights, based on the data of Rogers et al. [22]. Values for the stochastic algorithm are also from that publication. Models are arranged in order of decreasing bepe values.

| Model | Deterministic |  | Stochastic |  |
| --- | --- | --- | --- | --- |
|  | bepe | weight | bepe | weight |
| $\alpha$ | $1.13 \times 10^{-6}$ | 0 | $1.16 \times 10^{-6}$ | 0 |
| $\alpha\delta$ | $0.82 \times 10^{-6}$ | 0 | $0.87 \times 10^{-6}$ | 0 |
| $\alpha\gamma$ | $0.61 \times 10^{-6}$ | 0 | $0.62 \times 10^{-6}$ | 0 |
| $\alpha\gamma\delta$ | $0.40 \times 10^{-6}$ | 0 | $0.44 \times 10^{-6}$ | 0 |
| $\alpha\beta$ | $0.14 \times 10^{-6}$ | 0 | $0.18 \times 10^{-6}$ | 0 |
| $\alpha\beta\gamma$ | $0.14 \times 10^{-6}$ | 0 | $0.17 \times 10^{-6}$ | 0 |
| $\alpha\beta\delta$ | $0.11 \times 10^{-6}$ | 0.02 | $0.15 \times 10^{-6}$ | 0.16 |
| $\alpha\beta\gamma\delta$ | $0.10 \times 10^{-6}$ | 0.98 | $0.13 \times 10^{-6}$ | 0.84 |

When several models provide reasonable descriptions of the data, it is better to average across models than to choose just one. This allows uncertainty about the model itself to be incorporated into confidence intervals. For this purpose, Legofit uses bootstrap model averaging, “booma” [2, 20]. The booma weight of the *i*th model is the fraction of data sets (including the real data and 50 bootstrap replicates) in which that model “wins,” i.e. has the lowest value of bepe. The weights of all models are shown in table 3.

The new analysis, using the deterministic algorithm, replicates the main result of Rogers et al. [22]: that the most complex model (*αβγδ*) is preferred over all others. The strength of this preference, however, is stronger under the deterministic algorithm. The 2nd-place model (*αβδ*) gets 16% of the weight with the stochastic algorithm but only 2% with the deterministic one. The greater precision of the deterministic algorithm apparently improves Legofit’s ability to discriminate among models. The difference between these models is that *αβγδ* includes gene flow from early modern humans into Neanderthals, as proposed by Kuhlwilm et al. [12]. The current results strengthen the case for this hypothesis.

The model-averaged estimates of all parameters are shown in supplementary table S1. The two algorithms provide similar estimates, but there are two differences. First, the deterministic algorithm provides an unrealistic estimate of *T*_*XY*_, the separation time of Europeans and Africans.

This estimate—323 generations, or about 9000 y—is clearly too small. This may indicate that something is missing from the model or that identifiability problems have introduced bias. Further work would be needed to evaluate these alternatives. Second, the estimate of *N*_*S*_ is even larger—over 700,000—with the deterministic algorithm than with the stochastic one. This supports our previous suggestion that the superarchaic population was large or deeply subdivided [22].

## 4 Conclusions

Legofit’s new deterministic algorithm increases both speed and accuracy. The increase in accuracy results in smaller residual errors and better discrimination between alternative hypotheses. It has no large effect on confidence intervals, however, because these are primarily measuring uncertainty arising from statistical identifiability problems. The increase in speed is dramatic with models of small to moderate complexity and makes Legofit practicable on laptop computers. The deterministic algorithm slows dramatically, however, as models increase in complexity. For very complex models, the stochastic algorithm is still needed.

The deterministic algorithm replicated all the findings of Rogers et al. [22]. Because of its greater accuracy, it provided stronger support for the hypothesis that early modern humans contributed genes to Neanderthals [12]. It also strengthened the evidence that the superarchaic population was large or deeply subdivided [22].

Legofit is open source and freely available at https://github.com/alanrogers/legofit.

## Acknowledgements

I thank Greg Martin for comments on appendix B, Elizabeth Cashdan for editorial suggestions, and those who reviewed the manuscript for *PCI Mathematical and Computational Biology*. Analysis files are archived at doi:10.17605/OSF.IO/74BJF. This work was supported by NSF BCS 1638840, NSF BCS 1945782, and the Center for High Performance Computing at the University of Utah. Version 5 of this preprint has been peer-reviewed and recommended by *Peer Community In Mathematical and Computational Biology* (https://doi.org/10.24072/pci.mcb.100003).

## A The probability that *d* of *n* descendants derive from 1 of *k* ancestors

Eqn. 8 presents a formula for *Q*_*dk*_, the probability that a particular set of *d* descendants, chosen from a total of *n*, derives from a single unspecified ancestor, given that there were *k* ancestors in that ancestral generation. If *k* = 1, *Q*_*dk*_ = 1 as explained above. The result for *k >* 1 can be derived in two different ways.

### A.1 Short argument

Suppose that some ancestor has *d* descendants. The probability that a particular group of *d* descendents derives from this ancestor is 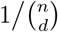, where 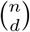 is the number of ways of choosing *d* descendants from a total of *n*. If *r* ancestors have *d* descendants each, the probability of descent from one of these is 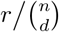. In reality, *r* is a random variable, and the probability becomes 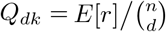, where *E*[*r*] is the expected value of *r*.

To derive *E*[*r*], number the ancestors from 1 to *k*, and let *y*_*i*_ represent the number of descendants of the *i*th ancestor, where *y*_*i*_ *>* 0 and Σ*y*_*i*_ = *n*. I will refer to a particular set of values, *y*_1_, …, *y*_*k*_, as an allocation of descendants among ancestors. The number of such allocations is 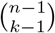 [6, pp. 38–39]. Furthermore, each allocation has the same probability, 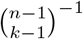, under the coalescent process [3, p. 13]. The *k* ancestors are statistically equivalent, which implies that 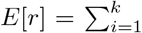 Pr{*y*_*i*_ = *d*} = *k*Pr{*y*_*i*_ = *d*} for an arbitrary ancestor *i*. If this ancestor has *d* descendants, there are 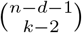 ways, each with probability 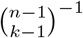, to allocate the *n*−*d* remaining descendants among the *k* − 1 remaining ancestors. Thus 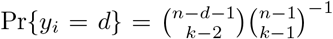,and *Q*_*dk*_ equals the expression in Eqn. 8.

### A.2 Longer argument

The *k* ancestors define a partition of the set of descendants into *k* subsets, each corresponding to a different ancestor. Let *x*_1_, *x*_2_, …, *x*_*k*_ denote the sizes of the *k* subsets, i.e., the numbers of descendants of the *k* ancestors. The probability of such a partition is given above in Eqn. 7. Suppose that a set of *d* descendants (and no others) derive from a single ancestor in interval *k*. This can happen only if *x*_*i*_ = *d* for some *i*. The ancestors are numbered in an arbitrary order, so let us set *x*_*k*_ = *d* and rewrite Eqn. 7 as

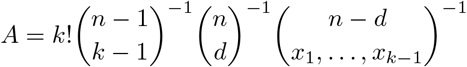

To calculate *Q*_*dk*_, we need to sum this quantity across all ways to partition the set of *n* − *d* remaining descendants into *k* − 1 subsets.

This is not the same as summing across values of *x*_*i*_, because each array of *x*_*i*_ values may correspond to numerous partitions of the set of descendants. This is illustrated in table 1, where the left side lists the 7 ways of partitioning a set of 4 descendants among 2 ancestors, along with the probability of each partition as given by Eqn. 7. The first four set partitions have equal probability, because each one divides the descendants into subsets of sizes 3 and 1, and the *x*_*j*_ values of these partitions therefore make equal contributions to Eqn. 7. Similarly, the last three set partitions have equal probability, because each divides the ancestors into two sets of size 2. These two cases: 3 + 1 = 4 and 2 + 2 = 4 are the two ways of expressing 4 as a sum of two positive integers. Eqn. 7 implies that all set partitions corresponding to a given integer partition have equal probability.

There are 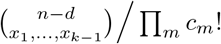 set partitions for a given partition of the integer *n*−*d* into *k* −1summands [1, theorem 13.2, p. 215]. In this expression, *c*_*m*_ is the number of times *m* appears among *x*_1_, …, *x*_*k*−1_. Multiplying this into *A* and summing gives

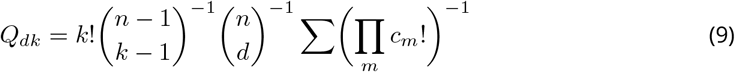

where the sum is over ways of partitioning *n* − *d* into *k* − 1 summands. Appendix B shows that this sum equals 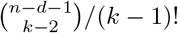. Substituting into Eqn. 9 reproduces Eqn. 8.

## B An identity involving integer partitions

The partition of a positive integer *n* into *k* parts can be written as 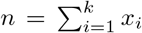, where the *x*_*i*_ are positive integers. This same partition is also *n* = Σ _*i*_ *ic*_*i*_, where *c*_*i*_ is the number of times *i* appears among the *x*_*i*_ values. In other words, *c*_*i*_ is the multiplicity of *i* in the partition. In terms of these multiplicities, *k* = Σ*c*_*i*_. This appendix will show that

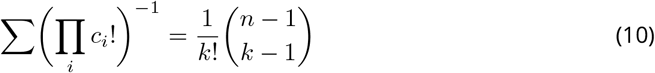

where the sum is across all partitions of an integer *n* into *k* parts.

This identity follows from the fact that there are 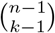 ways to put *n* balls into *k* boxes so that no box is empty [6, pp. 38–39]. Let us call each of these an “allocation” of balls to boxes. For each allocation, there is a corresponding partition of the integer *n* into *k* parts. The number of allocations often larger than the number of partitions. For example, there are 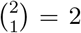 ways to put 3 balls into 2 boxes, **|* and *|**, where the stars represent balls and the bar separates boxes. Both allocations, however, correspond to a single partition, 3 = 2 + 1, of the integer 3. For a given integer partition, *c*_1_, *c*_2_, …, there are *k* !*/* П*c*_*i*_! distinct ways to allocate balls to boxes. (This is the number of ways to reorder the boxes while ignoring the order of boxes with equal numbers of balls.) The sum of this quantity across partitions must therefore equal 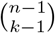. Dividing both sides by *k*! produces identity 10. Greg Martin posted a different proof of this identity on StackExchange.^2^

## Footnotes

1 https://github.com/alanrogers/legofit

2 https://math.stackexchange.com/questions/938280/on-multiplicity-representations-of-integer-partitions-of-fixed-length

## Notes

doi: 10.17605/OSF.IO/74BJF

### Competing Interest Statement

The authors have declared no competing interest.

### Summary of Updates

Version 5 of this preprint has been peer-reviewed and recommended by Peer Community In Mathematical and Computational Biology (https://doi.org/10.24072/pci.mcb.100003).

