## Supplementary methods for "An Efficient Algorithm for Estimating Population History from Genetic Data" (from another version of this preprint: version 4, 2021-05-03)

### Supplementary Materials*

Alan R. Rogers

May 3, 2021

#### S1 Simulations

In the archive, simulation code and simulated data sets are in directory `ae/sim`. To simulate 50 data sets using msprime [1], I executed the following bash command:

```
seq 0 49 | xargs -n 1 bash sim.sh
```

This invokes the shell script `sim.sh` 50 times. This shell script looks like this:

```
ofile=sim${1}.opf
efile=sim${1}.err
python3 msp.py -r | simpat 1>${ofile} 2>${efile}
```

Here, `simpat` is part of the Legofit package, and `msp.py` is a Python script that runs msprime. It is listed in appendix SA.1 on p. 3 and defines the model and all parameter values. This command generates 50 output files with names like `sim0.opf`, `sim1.opf`, and so on.

#### S2 Analysis of simulated data

##### S2.1 Deterministic algorithm

Analysis files using the deterministic algorithm can found in directory `ae/exact` within the archive. The pipeline in that directory is designed for use on a compute cluster running `slurm`, and the details of this pipeline can be found within that directory in the files `README.md`, `pipeline.sh`, `a1.slr` (appendix SB.2, p. 9), `a2.slr` (appendix SB.3, p. 9), `pclgo.slr` (appendix SB.4, p. 11), `b1.slr` (appendix SB.5, p. 11), and `b2.slr` (appendix SB.6, p. 12). However, this analysis is fast enough to run on an ordinary desktop computer, so I will describe the underlying Legofit commands here without reference to commands involving the cluster.

The analysis proceeds in several stages. Stages 1 and 2 use file `a.lgo`, which describes the simulation model accurately, except that the parameter values are perturbed away from their true values. Stage 1 executes the following command for data set 0:

```
legofit -1 -d 0 --stateOut a1-0.state --tol 3e-6 -S 5000 a.lgo \
  ../sim/sim0.opf > a1-0.legofit
```

*This supplement accompanies "An Efficient Algorithm for Estimating Population History from Genetic Data"

Similar commands are executed for each of the other 49 data sets. Here `a.lgo` is the input file describing the model of population history, `../sim/sim0.opf` is the simulated data set to be analyzed, and `a1-0.state` is the name of an output file that will record the optimizer's final state. The argument "`-1`" says not to ignore singleton site patterns, "`-d 0`" says to use the deterministic algorithm without ignoring states with low probability, "`--tol 3e-6`" tells the optimizer to stop when the spread of objective function values falls to $3 \times 10^{-6}$, and "`-S 5000`" tells the optimizer to do at most 5000 iterations. Legofit writes to standard output, which is redirected into `a1-0.legofit`.

Legofit uses the *differential evolution* optimization algorithm [2], which is quite good at finding global optima. Nonetheless, it is still possible that some of the 50 jobs in stage 1 will have gotten stuck on different local optima. To deal with this problem, each job in stage 2 of the analysis begins by reading the 50 `.state` files produced in stage 1, and generating an initial swarm of points that includes points from all 50 of the stage 1 jobs. Each stage 2 job is then able to choose among the optima found in stage 1. The legofit command for data set 0 in stage 2 looks like this:

```
legofit -1 -d 0 --tol 3e-6 -S 5000 a.lgo ../sim/sim0.opf \
  --stateIn a1-0.state \
  --stateIn a1-1.state \
  ...
  --stateIn a1-49.state > a2-0.legofit
```

This is just like the stage 1 command, except that there is no `--stateOut` command, and there are 50 `--stateIn` commands. Stage 2 generates 50 output files with names like `a2-0.legofit`.

Stages 1 and 2 present the optimizer with a challenge, because several of the free variables are tightly correlated, as shown in Fig. 3 of the main text. To alleviate this problem, stage 3 re-expresses the free variables in terms of principal components. To do this, `pclgo` reads `a.lgo` along with all 50 of the `.legofit` files produced in stage 3, and writes a single output file called `b.lgo`. The command that does this is

```
# (grep ^# a.lgo; pclgo a.lgo a2-*.legofit; grep -v ^# a.lgo |
 egrep -v "\<free\>") > b.lgo
```

This prints into `b.lgo` (1) the comments from `a.lgo`, (2) the free variables redefined as principal components, and (3) the rest of `a.lgo`. In previous publications, we have used the `--tol` argument of `pclgo` in order to reduce the number of dimensions. In the current analysis, we omit that argument, so there is no reduction in dimension. It is possible that reducing dimension can introduce bias, especially when there are identifiability problems in the data. Even without any reduction in dimension, `pclgo` makes the optimizer's job easier by re-expressing in orthogonal dimensions.

Stages 4 and 5 are just like stages 1 and 2, except that they use `b.lgo` instead of `a.lgo`.

##### S2.2 Stochastic algorithm

The stochastic algorithm is much slower and is best run on a compute cluster. The full pipeline, including slurm scripts, is available in the archive, but I will ignore cluster-related details here. Directory `ae/stoch` contains the code for and results from analysis under the stochastic algorithm, which follows the same steps as the deterministic one. The command for stage 1 looks like

```
legofit -1 --stateOut a1-0.state --tol 3e-5 \
  -S 5000@10000 -S 100@100000 -S 1000@2000000 a.lgo ../sim/sim0.opf \
  > a1-0.legofit
```

Note that the `-d 0` argument has been omitted. This tells `legofit` to use the default stochastic algorithm. I'm using a looser tolerance (`--tol 3e-5` rather than `--tol 3e-6`), because the stochastic algorithm can't achieve the same accuracy. Finally, the `-S` arguments are different. The first `-S` argument tells `legofit` to begin with 5000 generations of differential evolution, in each of which the objective function is evaluated using 10000 iterations of coalescent simulation. The next `-S` argument says to do 100 differential evolution generations with 100,000 coalescent iterations. The final one specifies 1000 differential evolution generations with 2,000,000 coalescent iterations. The optimizer will stop before these iterations are complete if it reaches the goal specified with `--tol`.

Stage 2 of the stochastic algorithm is similar to stage 2 of the deterministic one, except that it does not use the `-d 0` argument, and it specifies: `--tol 2e-5 -S 1000@2000000`. The tolerance is somewhat tighter than in stage 1, and there is only one `-S` argument.

Stage 3 of the stochastic algorithm is just like stage 3 of the deterministic algorithm. Stages 4 and 5 of the stochastic algorithm are just like stages 1 and 2, except that `b.lgo` is used instead of `a.lgo`.

### S3 Replication of results from Rogers et al. [4] using the deterministic algorithm

To illustrate the new deterministic algorithm in a realistic context, I reanalyzed the eight models considered by Rogers et al. [4], using the same data set, and the same bootstrap replicates. This analysis is described in section 2.8 of the main text. This supplement provides additional detail.

Each model is defined using an input file in .lgo format. The one for model $\alpha\beta\gamma\delta$ is in appendix SC.1, page 13. The 22 parameters defined there are listed in table S1.

The legofit program was run using the `-d 0` option, which invokes the deterministic algorithm, with a tolerance (`--tol 3e-6`) of $3 \times 10^{-6}$. This tolerance value is much smaller than that used by Rogers et al. [4]. It tells the estimator to attempt to fit the model to very high precision. One of the slurm scripts used in this analysis is shown in appendix SC.2 on page 14.

I used the bootstrap estimate of predictive error (bepe) for model selection and bootstrap model averaging (booma) for model averaging [3]. The model-averaged estimates of all parameters are shown in table S2.

### SA Simulations

#### SA.1 msp.py

```
#!/usr/bin/python3
import msprime
import os, sys, time

def usage():
    print("Usage: ./msp.py [options]")
    print("  where options may include:")
    print("  -r or --run: run simulation. Default: run")
    print("                DemographyDebugger")
    sys.exit(1)
```

Table S1: Parameters used in reanalysis of data from Rogers et al. [4]. The "text" column gives the symbol used in the text of this document; the "code" column gives the symbol used in input file `a.lgo` (appendix SC.1). Population sizes are in "haploid" units, which refer to twice the number of diploid individuals.

| Symbol | | |
|---|---|---|
| text | code | Description |
| $T_{XYND}$ | `Txynd` | separation time of $XY$ and $ND$ |
| $T_{XYNDS}$ | `Txynds` | separation time of $S$ |
| $T_{ND}$ | `Tnd` | separation time of $N$ and $D$ |
| $T_{N_0}$ | `Tav` | time of change in Neanderthal population size |
| $T_{XY}$ | `Txy` | separation time of $X$ and $Y$ |
| $T_D$ | `Td` | age of Denisovan fossil |
| $T_A$ | `Ta` | age of Altai Neanderthal fossil |
| $T_V$ | `Tv` | age of Vindija Neanderthal fossil |
| $T_\alpha$ | `TmN` | $\alpha$ admixture time |
| $T_\beta$ | `TmS` | $\beta$ admixture time |
| $T_\gamma$ | `TmXY` | $\gamma$ admixture time |
| $T_\delta$ | `TmSND` | $\delta$ admixture time |
| $2N_{N_0}$ | `twoNav` | size of early Neanderthal population |
| $2N_{N_1}$ | `twoNn` | size of late Neanderthal population |
| $2N_{ND}$ | `twoNnd` | size of population $ND$ |
| $2N_{XY}$ | `twoNxy` | size of population $XY$ |
| $2N_{XYND}$ | `twoNxynd` | size of population $XYND$ |
| $2N_S$ | `twoNs` | size of population $S$ |
| $m_\alpha$ | `mN` | $\alpha$ admixture fraction |
| $m_\beta$ | `mS` | $\beta$ admixture fraction |
| $m_\gamma$ | `mXY` | $\gamma$ admixture fraction |
| $m_\delta$ | `mSND` | $\delta$ admixture fraction |

Table S2: Parameter estimates and confidence intervals using real data and both algorithms. Results for stochastic algorithm are from Rogers et al. [4].

| Parameter | Deterministic | | Stochastic | |
|---|---|---|---|---|
| | Estimate | 95% C.I. | Estimate | 95% C.I. |
| $m_\alpha$ | 0.019 | (0.017, 0.021) | 0.019 | (0.018, 0.021) |
| $m_\beta$ | 0.023 | (0.021, 0.030) | 0.021 | (0.017, 0.027) |
| $m_\gamma$ | 0.009 | (0.008, 0.013) | 0.016 | (0.013, 0.025) |
| $m_\delta$ | 0.035 | (0.034, 0.052) | 0.034 | (0.022, 0.043) |
| $T_D$ | 3485 | (3394, 3729) | 3480 | (3242, 3665) |
| $T_V$ | 2471 | (2357, 2628) | 2506 | (2317, 2656) |
| $T_A$ | 5295 | (5208, 5452) | 5310 | (5078, 5415) |
| $T_{N_0}$ | 10644 | (10656, 12535) | 15706 | (14594, 16956) |
| $T_{ND}$ | 25135 | (25055, 25619) | 25400 | (25135, 25526) |
| $T_{XY}$ | 323 | (62, 506) | 1262 | (548, 2611) |
| $T_{XYNDS}$ | 78036 | (69103, 77977) | 79334 | (71901, 89765) |
| $2N_{N_1}$ | 5509 | (5532, 6278) | 6745 | (6444, 6994) |
| $2N_{N_0}$ | 15915 | (15192, 18389) | 31411 | (24767, 41586) |
| $2N_{ND}$ | 1618 | (597, 1756) | 1081 | (797, 1680) |
| $2N_{XY}$ | 58955 | (58716, 60979) | 55818 | (51085, 58416) |
| $2N_{XYND}$ | 40500 | (40214, 41453) | 40831 | (39917, 41264) |
| $2N_S$ | 1465235 | (557565, 2336867) | 53995 | (40181, 104336) |

```
do_simulation = False
for arg in sys.argv[1:]:
    if arg == "-r" or arg == "--run":
        do_simulation = True
    else:
        usage()

# time parameters in generations
Txynd = 25920
Tnd = 15000
Txy = 3788
Td = 1734      # age of Denisova fossil
Ta = 1760      # age of Altai fossil
Talpha = 1897     # time of Neanderthal admixture
Tepsilon = Td-1     # time of Denisovan admixture

# population sizes
Nxynd = 64964.1/2.0  # ancestral population
Nxy = 44869.2/2.0
Nnd = 5000/2.0
Nn = 9756.8/2.0
Nd = 5000/2.0
Nx = 20000/2.0 # modern Africa
```

```
Ny = 20000/2.0 # modern Europe

# admixture
mAlpha = 0.05
mEpsilon = 0.025

nchromosomes = 1000      # number of chromosomes
basepairs = 2e6   # number of nucleotides per chromosome
u_per_site = 1.4e-8 # mutation

# Recombination rate.  recomb is the probability of recombination
# between sites at opposite ends of the simulated sequence.
c = 1e-8 # rate per base pair per generation

# One haploid sample from each of 4 populations: two modern (X,Y),
# and two archaic (N,D).
samples = [
    msprime.Sample(population=0, time=0),  # population X
    msprime.Sample(population=1, time=0),  # population Y
    msprime.Sample(population=2, time=Ta), # population N
    msprime.Sample(population=3, time=Td)  # population D
]

lbl = ("x", "y", "n", "d")
npops = len(lbl)

# associate sample sizes with population labels
sampsize = {}
for s in lbl:
    sampsize[s] = 0

for samp in samples:
    ndx = samp.population
    assert ndx < len(lbl)
    assert lbl[ndx] in sampsize
    sampsize[lbl[ndx]] = 1

for s in lbl:
    if not (sampsize[s] > 0):
        print("Population %s has no samples" % s, file=sys.stderr)
        sys.exit(1)

# Population configurations. No sample sizes are listed here, because
# those are specified above in "samples".
popconf = [
    msprime.PopulationConfiguration(initial_size=Nx),
    msprime.PopulationConfiguration(initial_size=Ny),
    msprime.PopulationConfiguration(initial_size=Nn),
```

```
    msprime.PopulationConfiguration(initial_size=Nd)
]

events = [
    msprime.MassMigration(
        time=Tepsilon,
        source=1,
        dest=3,
        proportion=mEpsilon), # D->Y gene flow
    msprime.MassMigration(
        time=Talpha,
        source=1,
        dest=2,
        proportion=mAlpha), # N->Y gene flow
    msprime.MassMigration(
        time=Txy,
        source=1,
        dest=0,
        proportion=1.0), # X-Y split
    msprime.PopulationParametersChange(
        time=Txy,
        initial_size=Nxy,
        population_id=0),
    msprime.MassMigration(
        time=Tnd,
        source=3,
        dest=2,
        proportion=1.0), # N-D split
    msprime.PopulationParametersChange(
        time=Tnd,
        initial_size=Nnd,
        population_id=2),
    msprime.MassMigration(
        time=Txynd,
        source=2,
        dest=0,
        proportion=1.0), # XY-ND split
    msprime.PopulationParametersChange(
        time=Txynd,
        initial_size=Nxynd,
        population_id=0)
]

if do_simulation:
    # run simulation
    seed = int(time.time()) ^ os.getpid()
    sim = msprime.simulate(samples = samples,
                           population_configurations = popconf,
```

```
                              demographic_events = events,
                              length = basepairs,
                              recombination_rate = c,
                              mutation_rate = u_per_site,
                              num_replicates = nchromosomes,
                              random_seed = seed)

    # header
    print("npops = %d" % len(lbl))
    print("%s %s" % ("pop", "sampsize"))
    for s in lbl:
        print("%s %d" % (s, sampsize[s]))

    for i, chromosome in enumerate(sim):
        for variant in chromosome.variants():
            print(i, end=" ")
            for g in variant.genotypes:
                print(g, end=" ")
            print()

else:
    # Run demography debugger and quit
    dd = msprime.DemographyDebugger(
        population_configurations=popconf,
        demographic_events=events)
    dd.print_history()
    print("Use \"./sim -r\" to run simulation")
```

#### SB Analysis of simulated data using the deterministic algorithm

##### SB.1 Model of history: a.lgo

```
# Model: ((X,Y), (N,D)), migration: N -> Y, D -> Y
# This file is like true.lgo but the free parameter values have been
# arbitrarily moved to incorrect values so that legofit will have some
# work to do.
time fixed   zero  = 0
twoN fixed   one   = 1
time fixed   Txynd = 25920   # \citet[table~S12.2, p.~90]{Li:N-505-43-S88}
time free      Tnd = 20000
time free      Txy = 2500
time fixed     Talpha = 1897    # \citep[table~2]{Sankararaman:PLO-8-e1002947}
time fixed     Tepsilon = 1       # arbitrary
time free      Td  = 1200
time free      Ta  = 1100
twoN free   twoNnd = 800
twoN free    twoNn = 1000
```

```
twoN free    twoNd = 1000
twoN free   twoNxy = 20000
twoN free twoNxynd = 20000
mixFrac free    mAlpha = 0.01
mixFrac free    mEpsilon = 0.01
segment x    t=zero   twoN=one     samples=1
segment y    t=zero   twoN=one     samples=1
segment n    t=Ta     twoN=twoNn   samples=1
segment n2   t=Talpha    twoN=twoNn
segment d0   t=Tepsilon    twoN=one
segment d    t=Td     twoN=twoNd   samples=1
segment y1   t=Tepsilon    twoN=one
segment y2   t=Talpha    twoN=one
segment nd   t=Tnd    twoN=twoNnd
segment xy   t=Txy    twoN=twoNxy
segment xynd t=Txynd  twoN=twoNxynd
mix    y  from y1 + mEpsilon * d0
mix    y1 from y2 + mAlpha * n2
derive x  from xy
derive y2 from xy
derive n  from n2
derive n2 from nd
derive d0 from d
derive d  from nd
derive xy from xynd
derive nd from xynd
```

##### SB.2 Deterministic a1.slr

```
#!/bin/bash
#SBATCH -J a1
#SBATCH --account=rogersa-np
#SBATCH --partition=rogersa-np
#SBATCH --time=0:10:00
#SBATCH --nodes 1
#SBATCH --ntasks 1
#SBATCH -o a1-%a.legofit # stdout
#SBATCH -e a1-%a.err # stderr

i=${SLURM_ARRAY_TASK_ID}
ifile=‘printf "../sim/sim%d.opf" $i‘    # input file
stateout=‘printf "a1-%d.state" $i‘

time legofit -1 -d 0 --stateOut ${stateout} --tol 3e-6 -S 5000 a.lgo ${ifile}
```

##### SB.3 Deterministic a2.slr

```
#!/bin/bash
```

```
#SBATCH -J a2
#SBATCH --account=rogersa-np
#SBATCH --partition=rogersa-np
#SBATCH --time=0:10:00
#SBATCH --nodes 1
#SBATCH --ntasks 1
#SBATCH -o a2-%a.legofit # stdout
#SBATCH -e a2-%a.err # stderr

i=${SLURM_ARRAY_TASK_ID}
ifile=`printf "../sim/sim%d.opf" $i`    # input file

time legofit -1 -d 0 --tol 3e-6 -S 5000 a.lgo ${ifile} \
  --stateIn a1-0.state \
  --stateIn a1-1.state \
  --stateIn a1-10.state \
  --stateIn a1-11.state \
  --stateIn a1-12.state \
  --stateIn a1-13.state \
  --stateIn a1-14.state \
  --stateIn a1-15.state \
  --stateIn a1-16.state \
  --stateIn a1-17.state \
  --stateIn a1-18.state \
  --stateIn a1-19.state \
  --stateIn a1-2.state \
  --stateIn a1-20.state \
  --stateIn a1-21.state \
  --stateIn a1-22.state \
  --stateIn a1-23.state \
  --stateIn a1-24.state \
  --stateIn a1-25.state \
  --stateIn a1-26.state \
  --stateIn a1-27.state \
  --stateIn a1-28.state \
  --stateIn a1-29.state \
  --stateIn a1-3.state \
  --stateIn a1-30.state \
  --stateIn a1-31.state \
  --stateIn a1-32.state \
  --stateIn a1-33.state \
  --stateIn a1-34.state \
  --stateIn a1-35.state \
  --stateIn a1-36.state \
  --stateIn a1-37.state \
  --stateIn a1-38.state \
  --stateIn a1-39.state \
  --stateIn a1-4.state \
```

```
  --stateIn a1-40.state \
  --stateIn a1-41.state \
  --stateIn a1-42.state \
  --stateIn a1-43.state \
  --stateIn a1-44.state \
  --stateIn a1-45.state \
  --stateIn a1-46.state \
  --stateIn a1-47.state \
  --stateIn a1-48.state \
  --stateIn a1-49.state \
  --stateIn a1-5.state \
  --stateIn a1-6.state \
  --stateIn a1-7.state \
  --stateIn a1-8.state \
  --stateIn a1-9.state
```

##### SB.4 pclgo.slr

```
#!/bin/bash
#SBATCH -J pclgo
#SBATCH --account=rogersa-np
#SBATCH --partition=rogersa-np
#SBATCH --time=00:10:00
#SBATCH --nodes 1
#SBATCH --ntasks 1
#SBATCH -e pclgo.err # stderr

# Make b.lgo.
cmd="(grep ^# a.lgo;"
cmd+=" pclgo a.lgo a2-*.legofit;"
cmd+=" grep -v ^# a.lgo | egrep -v \"\<free\>\")"
echo "# "$cmd > b.lgo
eval $cmd >> b.lgo
```

##### SB.5 Deterministic b1.slr

```
#!/bin/bash
#SBATCH -J b1
#SBATCH --account=rogersa-np
#SBATCH --partition=rogersa-np
#SBATCH --time=0:10:00
#SBATCH --nodes 1
#SBATCH --ntasks 1
#SBATCH -o b1-%a.legofit # stdout
#SBATCH -e b1-%a.err # stderr

i=${SLURM_ARRAY_TASK_ID}
ifile=`printf "../sim/sim%d.opf" $i`    # input file
```

```
stateout=‘printf "b1-%d.state" $i‘

time legofit -1 -d 0 --stateOut ${stateout} --tol 3e-6 -S 5000 b.lgo ${ifile}
```

##### SB.6 Deterministic b2.slr

```
#!/bin/bash
#SBATCH -J b2
#SBATCH --account=rogersa-np
#SBATCH --partition=rogersa-np
#SBATCH --time=0:10:00
#SBATCH --nodes 1
#SBATCH --ntasks 1
#SBATCH -o b2-%a.legofit # stdout
#SBATCH -e b2-%a.err # stderr

i=${SLURM_ARRAY_TASK_ID}
ifile=‘printf "../sim/sim%d.opf" $i‘    # input file

time legofit -1 -d 0 --tol 3e-6 -S 5000 b.lgo ${ifile} \
  --stateIn b1-0.state \
  --stateIn b1-1.state \
  --stateIn b1-10.state \
  --stateIn b1-11.state \
  --stateIn b1-12.state \
  --stateIn b1-13.state \
  --stateIn b1-14.state \
  --stateIn b1-15.state \
  --stateIn b1-16.state \
  --stateIn b1-17.state \
  --stateIn b1-18.state \
  --stateIn b1-19.state \
  --stateIn b1-2.state \
  --stateIn b1-20.state \
  --stateIn b1-21.state \
  --stateIn b1-22.state \
  --stateIn b1-23.state \
  --stateIn b1-24.state \
  --stateIn b1-25.state \
  --stateIn b1-26.state \
  --stateIn b1-27.state \
  --stateIn b1-28.state \
  --stateIn b1-29.state \
  --stateIn b1-3.state \
  --stateIn b1-30.state \
  --stateIn b1-31.state \
  --stateIn b1-32.state \
  --stateIn b1-33.state \
```

```
  --stateIn b1-34.state \
  --stateIn b1-35.state \
  --stateIn b1-36.state \
  --stateIn b1-37.state \
  --stateIn b1-38.state \
  --stateIn b1-39.state \
  --stateIn b1-4.state \
  --stateIn b1-40.state \
  --stateIn b1-41.state \
  --stateIn b1-42.state \
  --stateIn b1-43.state \
  --stateIn b1-44.state \
  --stateIn b1-45.state \
  --stateIn b1-46.state \
  --stateIn b1-47.state \
  --stateIn b1-48.state \
  --stateIn b1-49.state \
  --stateIn b1-5.state \
  --stateIn b1-6.state \
  --stateIn b1-7.state \
  --stateIn b1-8.state \
  --stateIn b1-9.state
```

### SC Analysis of real data using the deterministic algorithm

#### SC.1 Model $\alpha\beta\gamma\delta$: a.lgo input file

```
# Gene flow: N->Y, S->D, XY->N, S->ND
# S = superarchaic; XY = early modern; ND = Neandersovan; Y = Europe;
# A, Altai; V=Vindija; D=Denisovan. Altai and Vindija on same branch
# TmN < Tv < Ta
time fixed  zero = 0
twoN fixed  one = 1
time fixed  TmN = 1        # no coalesc. events can happen b/t 0 and Tv
time fixed  Txynd = 25920  # \citet[table~S12.2, p.~90]{Li:N-505-43-S88}
time free   Txynds = 60000 # arbitrary: 1.74 myr
time free   Tnd = 24000
time free   Tav = 14000
time free   Txy = 10000
time free   Td  = 2500
time free   Ta  = 4000
time free   Tv  = 1000
twoN free   twoNav = 2000
twoN free   twoNn = 2000
twoN free   twoNnd = 2000
twoN free   twoNxy = 20000
twoN free   twoNxynd = 20000
twoN free   twoNs = 2000
```

```
mixFrac free   mN = 0.01
mixFrac free   mS = 0.01
mixFrac free   mSND = 0.01
mixFrac free   mXY = 0.01
time constrained TmXY = 0.5*(Txy + Tav)
time constrained TmS = 0.5*(Td + Tnd)
time constrained TmSND = 0.5*(Tnd + Txynd)
segment x    t=zero   twoN=one     samples=1
segment y    t=zero   twoN=one     samples=1
segment v    t=Tv     twoN=twoNn   samples=1
segment a    t=Ta     twoN=twoNn   samples=1
segment a2   t=TmXY   twoN=twoNn
segment d    t=Td     twoN=one     samples=1
segment d2   t=TmS    twoN=one
segment y2   t=TmN    twoN=one
segment n    t=TmN    twoN=twoNn
segment s    t=TmS    twoN=twoNs
segment s2   t=TmSND  twoN=twoNs
segment av   t=Tav    twoN=twoNav
segment nd   t=Tnd    twoN=twoNnd
segment nd2  t=TmSND  twoN=twoNnd
segment xy   t=Txy    twoN=twoNxy
segment xy2  t=TmXY   twoN=twoNxy
segment xynd t=Txynd  twoN=twoNxynd
segment xynds t=Txynds twoN=twoNxynd
mix    d    from d2 + mS * s
mix    y    from y2 + mN * n
mix    a    from a2 + mXY * xy2
mix    nd   from nd2 + mSND * s2
derive n    from v
derive v    from a
derive a2   from av
derive av   from nd
derive d2   from nd
derive x    from xy
derive y2   from xy
derive xy   from xy2
derive xy2  from xynd
derive nd2  from xynd
derive s    from s2
derive s2   from xynds
derive xynd from xynds
```

#### SC.2 a1boot.slr

```
#!/bin/bash
#SBATCH -J ABCDa1boot
```

```
#SBATCH --account=rogersa-kp
#SBATCH --partition=rogersa-kp
#SBATCH --time=36:00:00
#SBATCH --nodes 1
#SBATCH --ntasks 1
#SBATCH -o a1boot%a.legofit # stdout
#SBATCH -e a1boot%a.err # stderr

i=${SLURM_ARRAY_TASK_ID}
ifile=$(printf "../boot/boot%d.opf" $i)    # input file
stateout=$(printf "a1boot%d.state" $i)
lgofile=a.lgo

time legofit -1 -d 0 --stateOut $stateout --tol 3e-6 -S 5000 $lgofile $ifile
```
